# Divergent reward cue representations in prefrontal cortex underlie differences in reward motivation between adolescents and adults

**DOI:** 10.1101/2023.11.07.565069

**Authors:** Gabriela Manzano Nieves, Rachel M Rahn, Saba N Baskoylu, Conor M Liston

## Abstract

A prevailing view on postnatal brain development is that brain regions gradually acquire adult functions as they mature. The medial prefrontal cortex (mPFC) regulates reward learning, motivation, and behavioral inhibition, and undergoes a protracted postnatal maturation. During adolescence, reward-seeking behavior is heightened compared to adulthood — a developmental difference that may be driven by a hypoactive mPFC, with decreased top-down control of impulsive reward-seeking. However, this hypothesis has been difficult to test directly, due in part to technical challenges of recording neuronal activity *in vivo* across this developmental period. Here, using a novel 2-photon imaging-compatible platform for recording mPFC activity during an operant reward conditioning task beginning early in life, we show that the adolescent mPFC is hyper-responsive to reward cues. Distinct populations of mPFC neurons encode reward-predictive cues across development, but representations of no-reward cues and unrewarded outcomes are relatively muted in adolescence. Chemogenetic inhibition of GABAergic neurons decreased motivation in adolescence but not in adulthood. Together, our findings indicate that reward-related activity in the adolescent mPFC does not gradually increase across development. On the contrary, adolescent mPFC neurons are hyper-responsive to reward-related stimuli and encode reward-predictive cues and outcomes through qualitatively different mechanisms relative to the adult mPFC, opening avenues to developing distinct, developmentally informed strategies for modulating reward-seeking behavior in adolescence and adulthood.

## Introduction

Neuronal circuits undergo significant changes during postnatal development, which are thought to underpin age-specific behavioral variations observed in children and adolescents. In both humans and mice, adolescence is marked by significant increases in impulsivity, reward-seeking, and reward sensitivity compared to adults and pre-adolescents ^1–8^. Alterations in reward-seeking and sensitivity are also critical features of psychiatric disorders such as depression, obsessive-compulsive disorder, and attention deficit hyperactivity disorder, which often first emerge during adolescence ^9–11^. The medial prefrontal cortex (mPFC), one of the last brain regions to mature, plays a crucial role in modulating reward-seeking behavior and reducing impulsivity in adults ^12–17^. However, how the immature mPFC modulates reward-seeking behaviors in adolescence remains largely unknown. Elucidating how developmental changes in mPFC function impact reward processing across adolescence and adulthood is essential for understanding adolescent behavior and susceptibility to psychiatric conditions.

The mPFC continues to develop and mature through adolescence ^18–20^. mPFC glutamatergic (excitatory) ^21,22^ and GABAergic (inhibitory) neurons ^23,24^ undergo synaptic remodeling during adolescence, which likely results in different micro-connectivity in the adolescent mPFC compared to the adults. mPFC projections ^25^ and inputs to the mPFC continue to mature throughout adolescence ^7,18,27–29^, which can have profound effects on behavioral outcomes ^26^ and may influence how mPFC integrates information to guide behavior ^8,30^. That mPFC is still developing during adolescence is incontrovertible, but how these structural changes impact mPFC’s function and reward encoding properties remain unclear.

Pioneering neuroimaging studies using a variety of behavioral tasks have examined prefrontal cortical maturation across adolescence. For example, in go/no-go tasks, inhibitory control (no-go trial performance) increases from childhood into adulthood, ^31^ and is associated with increased volume of activation in the prefrontal cortex in children relative to adults ^32^. Other studies have revealed increased dorsolateral prefrontal cortex activation in adolescents ^33^ and children ^34^ compared to adults. However, some studies have reported the opposite effect, with increased dorsolateral prefrontal cortex activation in adults compared with children engaged in inhibitory control tasks, ^36^ positive correlations between age and prefrontal activity, ^35^ and mixed evidence of both increased and decreased PFC activity in adolescents compared to adults in a recent meta-analysis of fMRI studies ^37^. Thus, it is unclear whether PFC activity is increased or decreased in adolescents in various behavioral contexts based on fMRI measures alone. Here, we sought to test whether mPFC coding of reward-associated outcomes and reward-predictive cues increases across development from adolescence to adulthood by recording neuronal activity more directly, and whether the adolescent mPFC utilizes different coding mechanisms to represent these features.

Until recently, directly testing how the adolescent mPFC represents reward signals has been technologically difficult. Recording neuronal activity from growing animals while they learn and perform a task presents various challenges, as mice grow in size and traverse adolescence within 20 days. To overcome methodological difficulties, we created custom-made behavioral apparatuses and adapted prism implant techniques suitable for adolescent mice, allowing us to record neuronal calcium activity via an implanted optical prism ^38–40^ across development. We used a custom reward-association behavioral task that allowed animals to be trained and tested during adolescence. We leveraged these innovations to test if the adolescent mPFC is recruited during reward tasks, and if so, if it exhibits changes in neuronal representation and function.

## Results

### Adolescents have increased reward acquisition and drive compared to adults

To assess reward-motivated behaviors, we developed an active reward task compatible with head-fixed 2-photon imaging in both adults (>60 days old) and adolescents (28 - 45 days old). The active reward task permits analysis of general learning, motivational drive, and inhibitory control, and is akin to a go no-go task. A trial begins with the onset of a conditioned stimulus (CS, tone, 5-second duration), during which a mouse’s response (lick to spout) determines the trial outcome (**Fig. 1a**). If the reward-predicting CS (CS+) plays, mice can lick the spout (Hit) to obtain a sucrose reward at tone offset, or not lick the spout (miss) and forgo a reward. For the first 3 days, mice are trained to 100 trials of a single rewarded tone (CS+ only, **Fig. 1b**), which allows mice to learn the CS+ association while avoiding overtraining. Following CS+ only training, a CS-that is never rewarded is introduced (**Fig. 1c**) to allow us to compare behavioral responses between reward and no-reward cues (50 trials of each CS). If mice lick to the CS-, it is considered a False Alarm; abstaining from licking during the CS- tone is a correct rejection.

**Figure 1:**
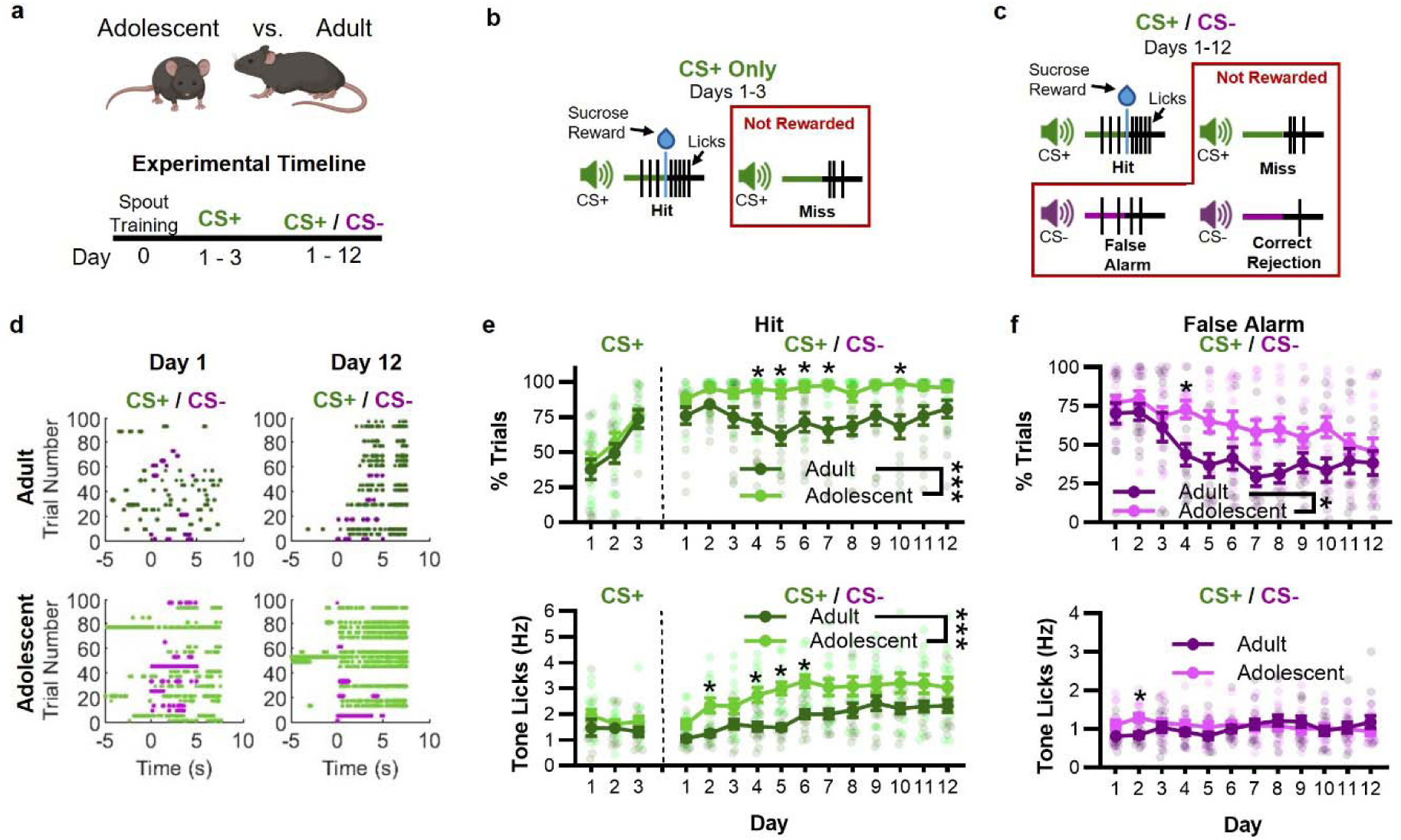
Adolescent mice have increased reward motivation. a) Experimental timeline. b) Schematic of CS+ only training sessions. c) Schematic of CS+/CS- training session. d) Rasterized lick times for an adult (top row) and adolescent (bottom row) mouse during early (day 1) and late (day 12) CS+/CS- training sessions. Plots are aligned to tone onset (at 0 s). If rewarded, a sucrose droplet is delivered at tone offset (at 5 s). CS+ trials are green, CS- trials are magenta. Every 4th trial (25 trials) is shown for ease of visualization. e) Graphs of % acquired CS+ trials (top) and CS+ lick rates during Hit trials (bottom). Adolescent mice have significantly higher % trials (F_1,26_ = 21.70, p < 0.0001), and tone lick rate during Hit trials (F_1,26_ = 15.27, p = 0.0006) compared to adults. Mice generally increase tone lick rates across training days (F_3.8,99.7_ = 12.11, p < 0.0001). Data is shown as mean ± SEM. A two-way repeated measures ANOVA was used for the main effects. * indicates p < 0.05 on Šídák’s multiple comparison post hoc test. n = 10-15 mice per age group. f) Graphs of % acquired CS- trials (top) and CS- lick rates in False Alarm trials (bottom). Adolescent mice obtained significantly higher False Alarm trials (CS- trials where they lick) (F_1,26_ = 7.16, p = 0.012), but not False Alarm lick rates (F_1,26_ = 0.49, p = 0.48), compared to adults. Mice generally decreased % False Alarm trials acquired across training days (F_4.6,121.5_ = 9.59, p < 0.0001). A significant interaction of age by training day was observed (F_11,284_ = 2.15, p = 0.016) for False Alarm tone licks. Data is shown as mean ± SEM. Main effects were determined with a two-way repeated measures ANOVA (top), or mixed-effects model analysis (bottom). * indicates p < 0.05 on Šídák’s multiple comparison post hoc test. n = 10-15 mice per age group.

To test behavioral differences between adolescent and adult mice, we trained and tested two separate age cohorts of mice on the active reward task (**Fig. 1d**). We found that adolescents and adults learned simple associations (CS+ only) at a similar rate (**Fig. 1e**), suggesting both groups can learn simple associations. However, with the addition of the CS- trials, adolescent and adult behavioral responses quickly diverged. Adolescents had more Hit trials and a higher lick rate to Hit trials compared to adults (**Fig. 1e**), consistent with a greater motivational drive. Adolescent mice also had more False Alarm outcomes than adults (**Fig. 1f**). The increased False Alarms in adolescents suggest diminished inhibitory control. Similar to adults, adolescents displayed a lower lick rate to False Alarm trials compared to Hit trials (**Extended Data Fig. 1**), suggesting adolescents do understand that the CS- is not rewarded. Together, our data corroborates previous findings that adolescent mice have increased reward-seeking behavior.

### Adolescent neuronal representation displays increased activity on Hit trials

Previous work indicates that the mPFC undergoes a protracted development, and that the adolescent mPFC may be hypoactive in a variety of behavioral contexts compared to the adult mPFC. To test this, we used two-photon calcium imaging to determine whether representations of rewarded trials and reward-predictive cues within the adolescent mPFC are diminished compared to adults. We hypothesized that diminished representation of reward-related information might be observed as either a reduction in the proportion of neurons encoding task-relevant information or a decrease in magnitude of reward-related activity in the adolescent mPFC compared to adults. To test these hypotheses, we used chronically implanted microprisms and 2-photon calcium imaging to record mPFC neuronal activity as mice performed the active reward task. To simultaneously record from glutamatergic and GABAergic mPFC neurons, we used a viral vector to express the calcium indicator GCaMP8f pan-neuronally in transgenic mice that expressed a red reporter (Ai9) in GABAergic neurons (Gad2-cre) (**Fig. 2a**). This strategy allowed us to distinguish glutamatergic (Ai9 negative) and GABAergic (Ai9 positive) neurons (**Fig. 2b**). To quantify neuronal activity related to reward and no-reward cues, we performed two-photon calcium imaging on day 7 (**Fig. 2c**) after mice had learned the CS+/CS- associations. To investigate the coding properties of mPFC neurons, we used a general linear model (GLM) with three predictors representing the most important features of the task (CS+, CS-, and Lick) to identify task-relevant neurons (**Fig. 2d**; see Online Methods for details).

**Figure 2:**
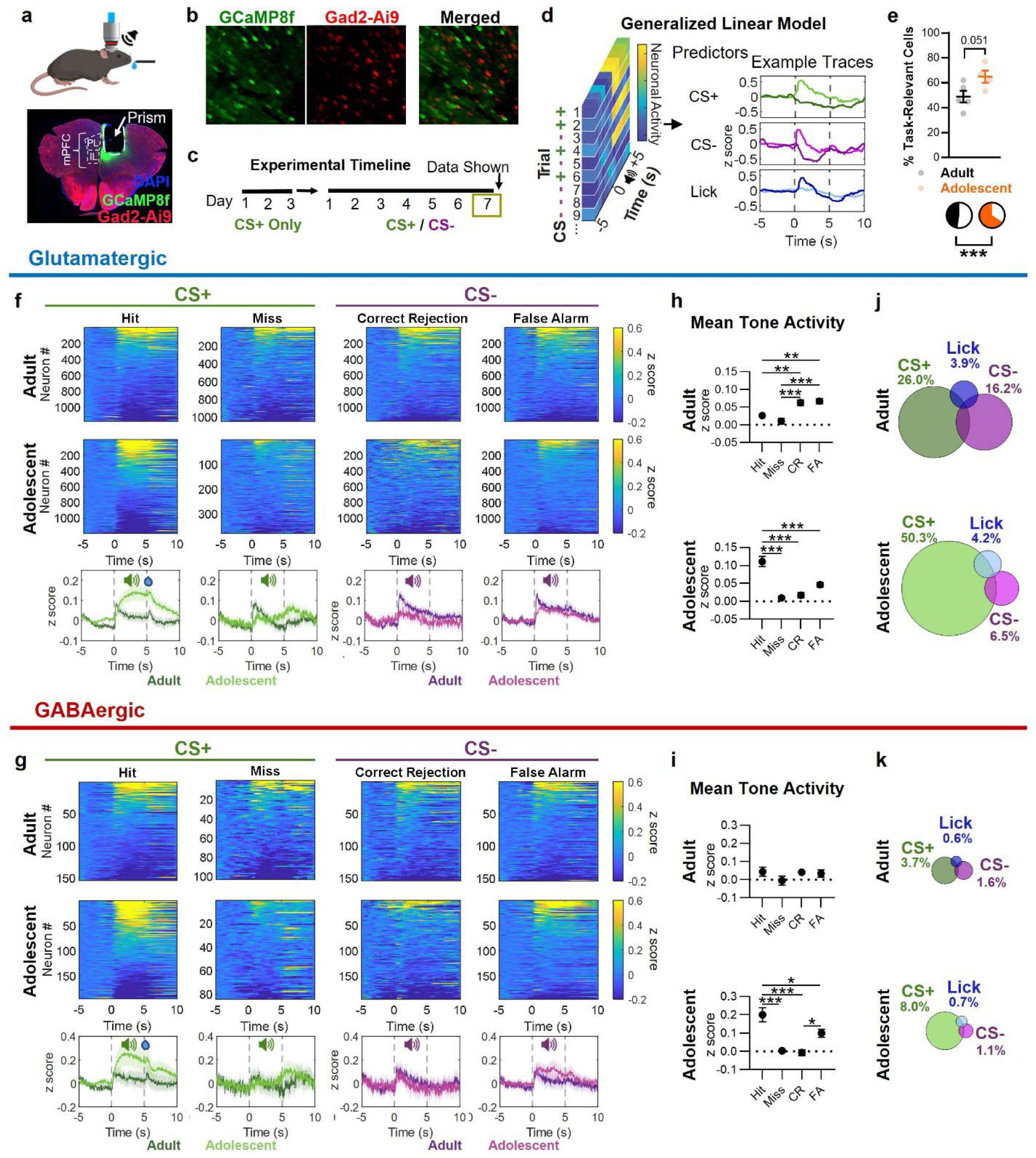
Adolescent mPFC overrepresents reward cues and exhibits increased reward cue activity. a) Illustration of a mouse during 2P imaging and histological image of prism implant in mPFC. b) Representative images of glutamatergic (green only) and GABAergic (co-labeled green and red) neurons during 2P imaging. Images were captured using Galvano imaging and processed using a max intensity projection. Neurons were recorded from 5 adult and 4 adolescent mice. c) Experimental timeline of behavioral training. 2P imaging data was extracted from day 7 of CS+/CS- training. d) Schematic of generalized linear model (GLM) used to classify neurons. 3 predictors were used: CS+, CS-, and Lick. Pre-tone and Tone activity (-5 to 5 seconds) was used in the GLM. e) Plot of individual values and mean percent of task-relevant neurons in adult and adolescent mice (top) (t.test: t 7 = 2.34, p = 0.051), and pie charts of pooled neurons from all task-relevant (colored) and non-task-relevant (white) neurons (Chi^2^: χ^2^_(1)_ = 159.5, p < 0.0001). f) Heatmap of neuronal activity of task predictive glutamatergic neurons. Neurons are sorted from greatest to least mean tone activity (0 to 5 seconds). Neuronal activity of adults (top), adolescents (middle), and the average neuronal trace (bottom; average of heatmaps) are presented. g) Heatmap of neuronal activity of task predictive GABAergic neurons, displayed as in Fig. 2g. h) Plot of glutamatergic neuron’s mean tone activity (0 to 5 seconds) for each outcome in adults (top) and adolescents (bottom). A main effect of outcome was found in adults (F_3, 4594_ = 11.81, p < 0.0001), and adolescents (F_3, 3969_ = 19.84, p < 0.0001). One-way ANOVA followed by Tukey’s multiple comparison analysis. Data is shown as mean ± SEM. * p < 0.05, ** p < 0.01, *** p < 0.001 i) Plot of GABAergic neurons mean tone activity (0 to 5 seconds) for each outcome for adults (top) and adolescents (bottom). A main effect of outcome was found in adolescents (bottom; F_3, 668_ = 12.69, p < 0.0001), but not adults. One-way ANOVA followed by Tukey’s multiple comparison analysis. Data is shown as mean ± SEM. * p < 0.05, ** p < 0.01, *** p < 0.001 j) Venn diagram of glutamatergic mPFC neurons significant for GLM predictors (CS+, CS- and/or Lick), comparing adult (top), and adolescent (bottom) neurons (Chi^2^: χ^2^_(2)_ = 232.2, p < 0.0001). The percent of neurons are shown. For n of neurons see **Extended Data Fig. 3**. k) Venn diagram of GABAergic mPFC neurons significant for GLM predictors (CS+, CS- and/or Lick), comparing adult (top), and adolescent (bottom) neurons (Chi^2^: χ^2^_(2)_ = 19.47, p < 0.0001). The percent of neurons are shown. For n of neurons see **Extended Data Fig. 3**.

First, we quantified the proportion of neurons with task-related activity, defined as neurons modulated by one or more task features. We found that a significantly larger proportion of adolescent neurons encoded one or more task features compared to adults (**Fig. 2e**). In adults, we found that 1335 out of 2795 neurons (47.7%) were significantly task-relevant, with 154 of the task-relevant neurons being GABAergic. In adolescence, 1394 out of 2117 recorded neurons (65.8%) were found to be task-relevant, with 196 of the task-relevant neurons being identified as GABAergic neurons.

To directly test how adult and adolescent task-relevant neurons respond to the four possible outcomes (Hit, Miss, Correct Rejection, and False Alarm), we compared the mean tone activity (0-5 s tone) for each of the four trial outcomes for glutamatergic (**Fig. 2f**) and GABAergic (**Fig. 2g**) neurons. In adult glutamatergic neurons (**Fig. 2h top**), the mean activity of all task-relevant neurons was significantly greater during correct rejections and False Alarms (both CS- outcomes) compared to Hit and miss outcomes (both CS+ outcomes). In contrast, in adolescence, we found that glutamatergic neurons had a significantly greater mean activity during Hit outcomes compared to each of the other outcomes (**Fig. 2h bottom**), and when compared to Hit trials in adults (**Extended Data Fig. 2**). This suggests that activity in the adolescent mPFC is dominated by reward cue responses, which are amplified compared to adults, while the adult mPFC is most active on non-rewarded trials.

Next, we repeated this analysis for GABAergic neurons (**Fig. 2i**). As above, in adolescents, mean GABAergic neuron activity was increased during Hit trials (rewarded CS+ tones) compared to other trial outcomes (**Fig. 2i bottom**), while in adults, we did not find any mean tone activity differences across the four task outcomes (**Fig. 2i top**). Like adolescent glutamatergic neurons, adolescent GABAergic neurons also demonstrated increased mean tone activity during Hits compared to adults (**Extended Data Fig. 2**). Together, these data suggest that adolescent glutamatergic and GABAergic neurons are more responsive to rewarded (Hit) trials compared to adults, indicating that the adolescent mPFC is more attuned to reward information.

To determine if age influenced the task feature that neurons encoded, we analyzed the proportion of all glutamatergic (**Fig. 2j**) or GABAergic (**Fig. 2k**) neurons with activity significantly modulated by each task predictor (CS+, CS-, Lick). We found that adolescents had a larger percentage of CS+ encoding neurons in both glutamatergic (50.3% vs. 26% in adults) and GABAergic (8% vs. 3.7% in adults) cell populations. Inversely, adults had a larger percentage of CS- encoding neurons in both glutamatergic (16.2% vs. 6.5% in adolescents) and GABAergic (1.6% vs. 1.1% in adolescents) cell populations. Together, these data indicate that most task-modulated cells encode the reward-predictive (CS+) cue. In adolescence however, CS+ encoding cells are *over*-represented while CS- encoding cells *under*-represented compared to adults.

### Adolescent mPFC is tuned to reward predictive information

Upon identifying functional subtypes of neurons whose activity is modulated by specific task features, we next sought to test whether neuronal activity in the mPFC is sufficient to encode reward outcomes and the identity of reward-predictive cues, and whether these coding properties differed in adolescents vs. adults. To this end, we trained support vector machine (SVM) classifiers to decode reward outcomes or CS identity and tested whether their performance varied by age or by functional cell type. SVM decoders were trained on tone activity for all mPFC neurons or for specific functional subtypes (e.g. CS+, CS-, lick) in 70% of trials, and performance was evaluated in the remaining 30% across 20 iterations of cross-validation (**Fig. 3a**; see Online Methods). We found that reward outcomes could be decoded with high accuracy rates from activity in all functional subtypes, but accuracy rates were significantly higher for adolescents than for adults (**Fig. 3b**). A similar pattern was observed for decoding CS identity, but performance differences for adolescents vs. adults were smaller (**Fig. 3c**).

**Figure 3:**
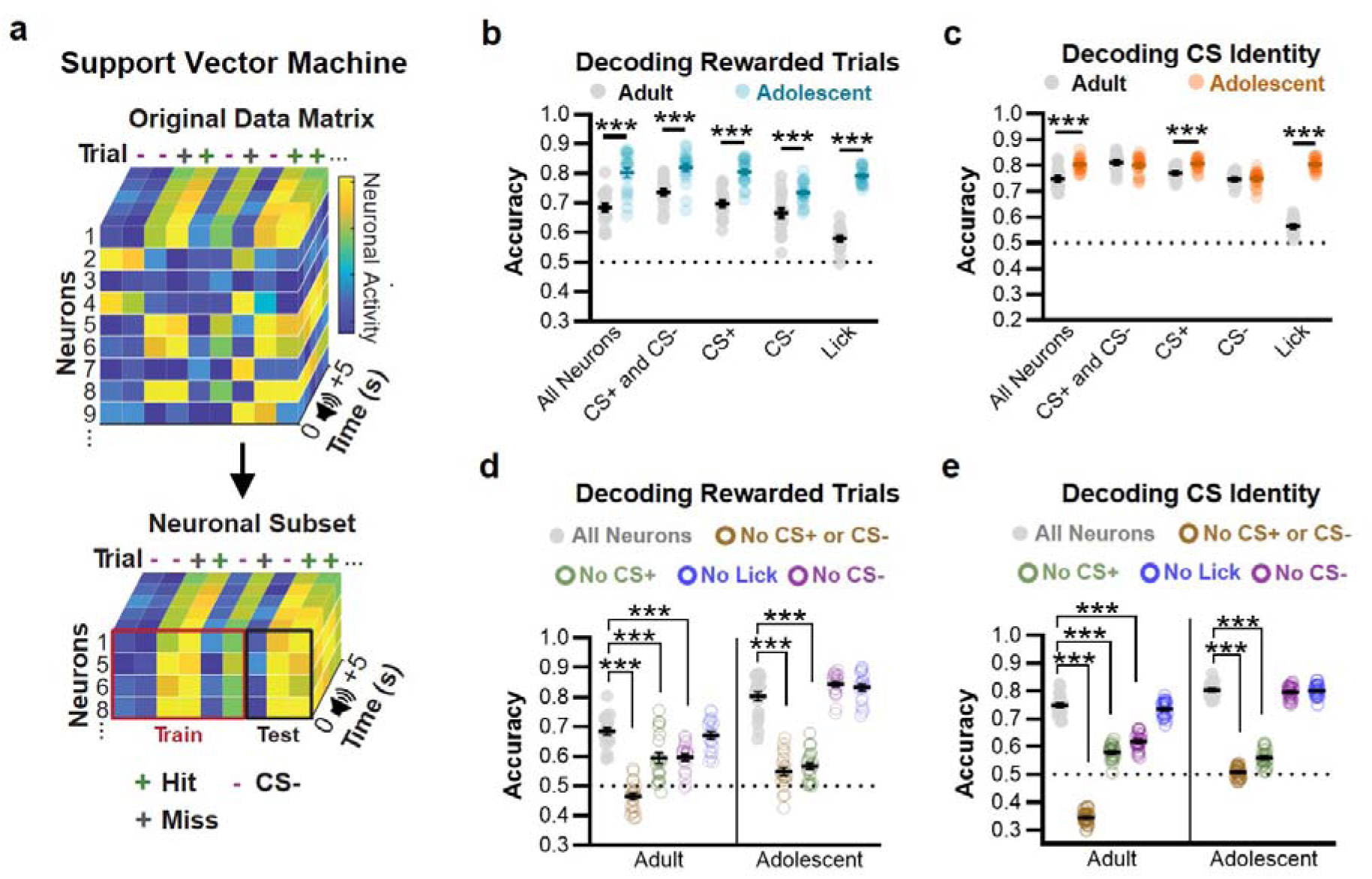
Adolescent mPFC has more cue and reward predictive information than the adult mPFC. a) Schematic of Support Vector Machine (SVM) decoding of neuronal subpopulations (left). SVMs were run on matrices containing glutamatergic and GABAergic neurons. The 5 seconds of tone activity was used for decoding. 20 train/test sessions were run for each subpopulation. For each run, the 120 CS trials were randomly shuffled, with the time series remaining intact. For comparison against shuffled data, within age effects, and full statistics see **Extended Data Fig. 4**. b) Test of the sufficiency of neuronal subsets to decode Rewarded trials (Hit versus Miss, Correct Rejection, and False Alarm). X-axis labels indicate neuronal subsets included in the SVM decoder model. Individual trials and mean ± SEM for the decoder is shown. A Tukey multiple comparison analysis was used. * p < 0.05, ** p < 0.01, *** p < 0.001. c) Test of the sufficiency of neuronal subsets to decode CS identity (CS+ versus CS- trials). X-axis labels indicate neuronal subsets included in the SVM decoder model. Individual trials and mean ± SEM for the decoder is shown. A Tukey multiple comparison analysis was used. * p < 0.05, ** p < 0.01, *** p < 0.001. d) Test of the necessity of neuronal subsets to decode Rewarded trials (Hit versus Miss, Correct Rejection, and False Alarm). Individual trials and mean ± SEM for the decoder is shown. A Tukey multiple comparison analysis was used. * p < 0.05, ** p < 0.01, *** p < 0.001. e) Test of the necessity of neuronal subsets to decode CS identity (CS+ versus CS- trials). Individual trials and mean ± SEM for the decoder is shown. A Tukey multiple comparison analysis was used. * p < 0.05, ** p < 0.01, *** p < 0.001.

To further understand whether particular functional subtypes of mPFC neurons were necessary for encoding reward outcomes or CS identity, we examined how removal of each neuronal subset from the larger neuronal population affected decoding accuracy. In adults, removal of either CS+ neurons, CS- neurons, or both decreased the decoding accuracy for reward outcomes (**Fig. 3d**) and CS identity (**Fig. 3e**). In contrast, in adolescents, removing CS+ neurons decreased decoding accuracy to near chance levels for both CS identity and reward outcomes, but removing CS- neurons had no effect (**Fig. 3d-e**). Together, these results indicate that the adolescent mPFC encodes representations of reward-predictive cues that are at least as strong as those encoded by the adult mPFC, and that the adolescent mPFC significantly outperforms the adult mPFC in its ability to encode reward outcomes. During adolescence, the mPFC relies primarily on CS+ neurons to encode reward outcomes and CS identity, whereas in adulthood, these representations are distributed across multiple functional subtypes.

### Adolescent mPFC is tuned to reward predictive information

To understand how different neuronal subpopulations encode information in adults and adolescents, we next analyzed the changes in activity patterns in CS+, CS-, and Lick encoding neurons by trial outcome (**Fig. 4a-b**). To visualize how activity patterns in functional subtypes of glutamatergic mPFC neurons vary with age, we identified groups of neurons that were significantly activated or inhibited by each task feature (i.e. significant positive or negative beta weights for the CS+, CS–, or Lick predictors in the GLM depicted in **Fig. 2**) and plotted their mean activity by trial outcome (Hit, Miss, Correct Rejection, False Alarm). In adults, we found that mean tone activity in CS+ encoding neurons was significantly different during Hit trials compared to either of the two CS- outcomes (Correct Rejection, False Alarms), but not between Hit and Miss outcomes (**Fig. 4c**). In contrast, in adolescents, mean tone activity in CS+ encoding neurons was significantly different during Hit trials compared to all unrewarded outcomes, including Miss trials (**Fig. 4c**). This provides additional evidence that adolescent CS+ encoding neurons are strongly tuned to whether a trial was rewarded.

**Figure 4:**
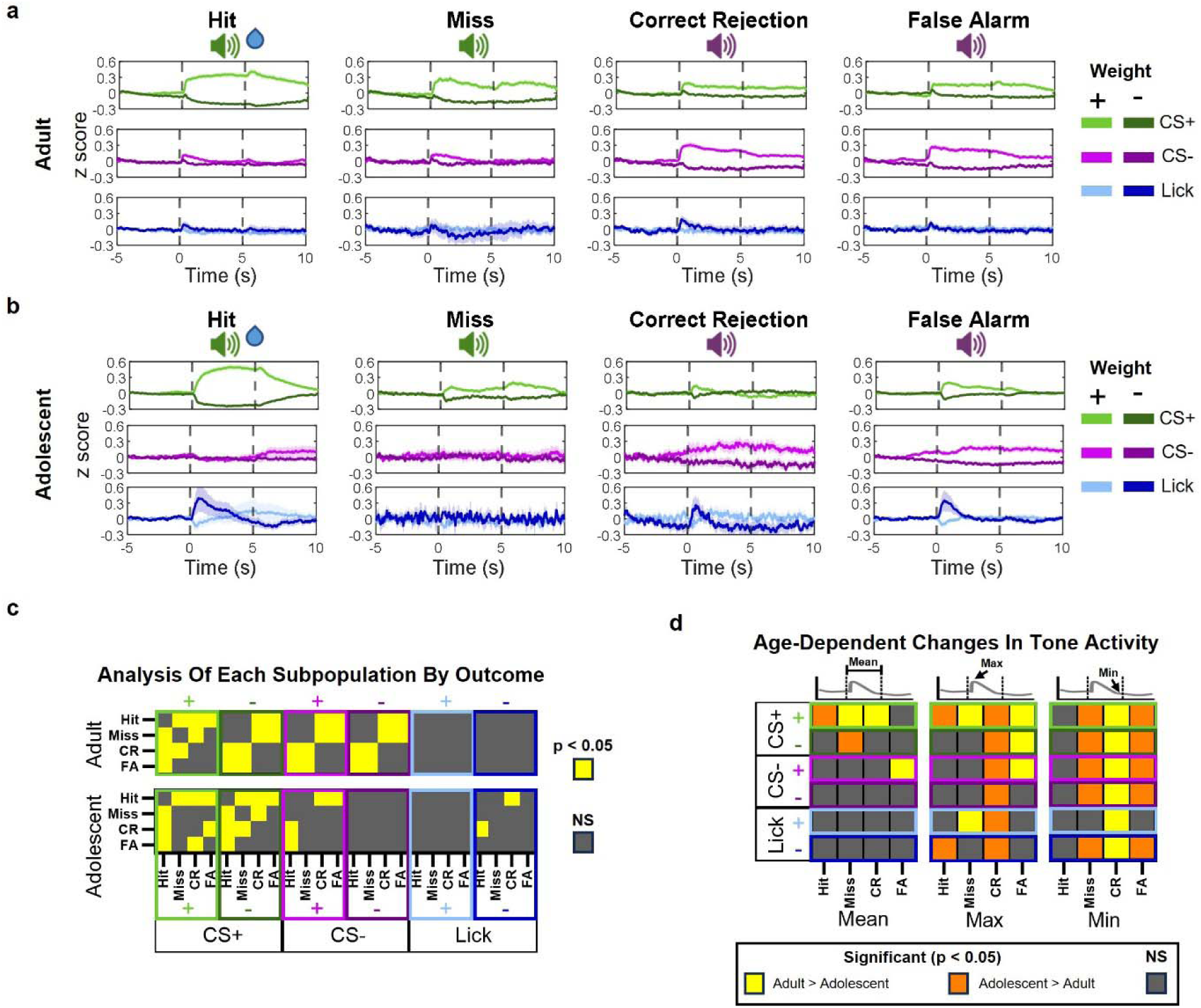
Neuronal subpopulation activity changes across outcomes. a) Mean activity for every subpopulation of adult neurons during each outcome. For each predictor, two lines are shown: a negative beta weight predictor (darker line) and a positive predictor (lighter line). Graphs show mean ± SEM (shaded area). b) Mean activity for every subpopulation of adolescent neurons during each outcome. For each predictor, two lines are shown: a negative beta weight predictor (darker line) and a positive predictor (lighter line). Graphs show mean ± SEM (shaded area). c) Within-age analysis of subpopulations of neurons in the adult (top) and adolescent (bottom) mPFC. For analysis, a two-way ANOVA followed by a Tukey multiple comparison test was used to determine if the mean tone activity of each subpopulation significantly changed across outcomes (Hit, Miss, Correct Rejection, False Alarm). The heatmap shows the significant (p < 0.05) and non-significant Tukey post-hoc comparisons. See **Extended Data Fig. 5** for full statistical analysis. d) Heatmap of significant comparisons between the adult and adolescent mean (left), maximum (middle), and minimum (right) tone activity for each neuronal subpopulation. A separate two-way ANOVA, containing tone data for all 6 neuronal subpopulations, was run for each of the four outcomes. The heatmap shows the significant (p < 0.05) and non-significant Šídák post-hoc comparisons of each subpopulation. See **Extended Data Fig. 6** for full statistical analysis.

CS- encoding neurons exhibited a strikingly different pattern. Adult CS- encoding neurons had significantly different mean tone activity between CS+ trial outcomes and CS- trial outcomes, whereas adolescent CS- encoding neurons did not (**Fig. 4c and Extended Data Fig. 5**). Furthermore, analysis of age-related differences in the mean, maximum, and minimum of the tone activity for each subpopulation revealed that the neuronal subpopulations in general respond similarly across ages when the trial is a Hit compared to all other outcomes (**Fig. 4d and Extended Data Fig. 6**). This further suggests that the main differences in age may not be in the encoding of the reward but in the encoding of the differences between the unrewarded vs rewarded outcomes.

### Adolescent mPFC has an expanded role in reward-seeking

Having identified the role of mPFC neurons in the encoding and decoding of reward cues, we sought to next test the causal contribution of glutamatergic (VGlut1-cre) and GABAergic (Gad2-cre) neurons to adolescent and adult reward-seeking behavior. We had previously found that GABAergic adult neurons did not significantly change their mean tone activity dependent on the outcome, but adolescent GABAergic neurons did (**Fig 2k**). Thus, we hypothesized that adolescent GABAergic neurons would have a more profound impact on behavior than adult GABAergic neurons. To test if the adolescent mPFC was more capable of driving reward-seeking behavior than the adult mPFC, we used chemogenetics to excite (hM3D(Gq)) or inhibit (hM4D(Gi)) neurons in the mPFC (**Fig. 5a and Fig. 5c**). All groups (control, inhibition, and excitation) were administered a chemogenic agonist CNO on days 6 and 7 of training (**Fig. 5b**). This timeline allows mice the necessary 5 days to learn the difference between CS+ and CS- tones while avoiding overtraining which could mask mPFC’s role in the task. Data is expressed as a change in behavior from day 5 of training (equation shown in **Fig. 5b**).

**Figure 5:**
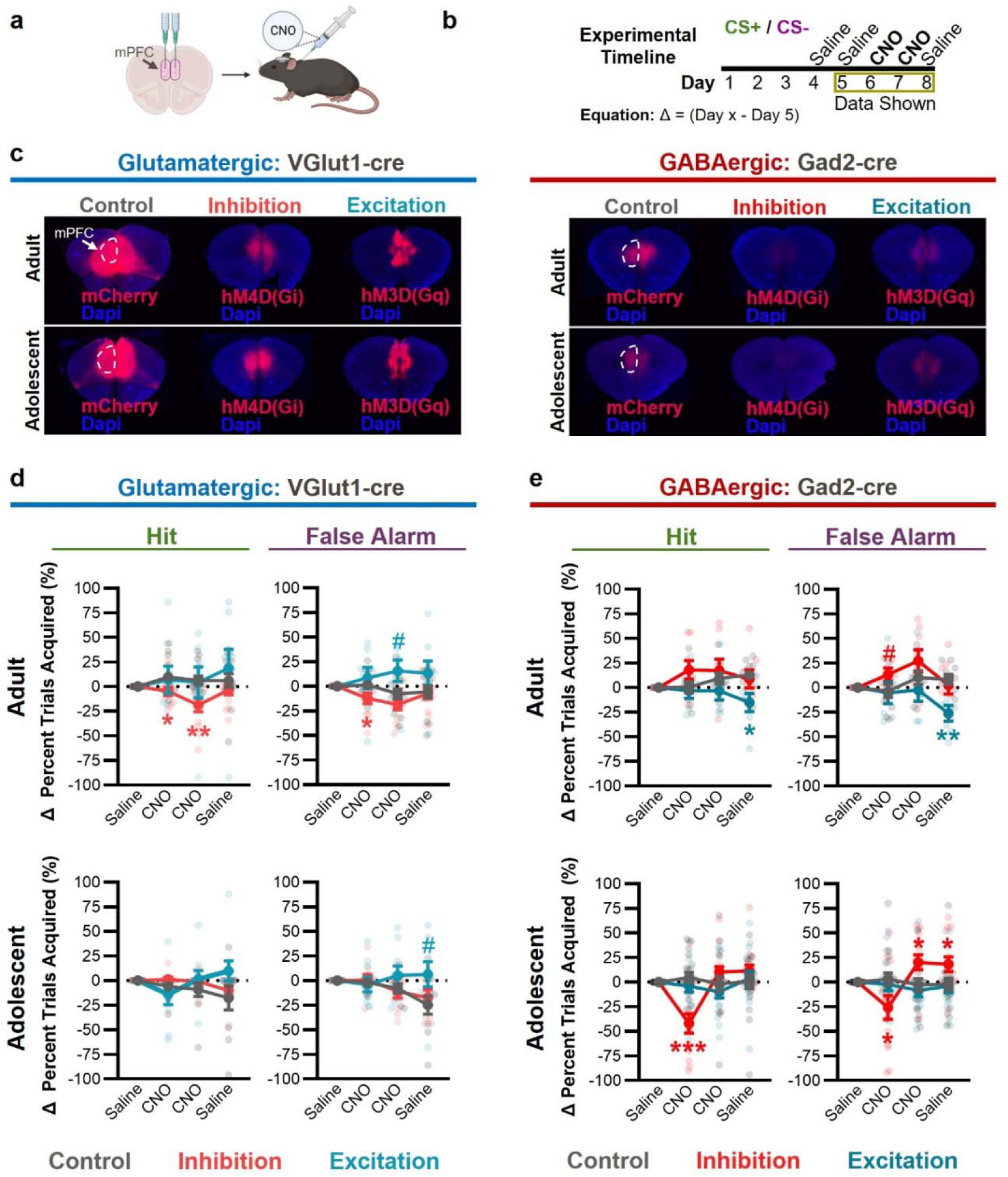
Adolescent mPFC has an expanded role in reward-seeking. a) Illustration of viral injections and CNO administration in mice. b) Experimental timeline. All mice are injected with saline solution on days 4 and 5, CNO on days 6 and 7, and saline on day 8. The equation used to measure the change is displayed below. c) Pictures of DREADD viral expression in glutamatergic (left) and GABAergic (right) neurons. d) Inhibition (red) and excitation (blue) of glutamatergic mPFC neurons in adult (top) and adolescent mice. A Tukey’s multiple comparison analysis was run per graph. n = 8-14 mice per condition per age. # p <0.10, * p < 0.05, ** p < 0.01, *** p < 0.001 e) Inhibition (red) and excitation (blue) of GABAergic mPFC neurons in adult (top) and adolescent mice. A Tukey’s multiple comparison analysis was run per graph. n = 5-19 mice per condition per age. # p <0.10, * p < 0.05, ** p < 0.01, *** p < 0.001. For 3-way ANOVA analysis of age by treatment by day effects see **Extended Data Table. 1**

Modulation of glutamatergic and GABAergic neurons revealed age-specific differences in activity. Inhibiting adult glutamatergic neurons decreased the percent of Hit and False Alarm trials (**Fig. 5d top**), whereas inhibiting glutamatergic neurons in adolescents had no effect on behavior (**Fig. 5d bottom**). Notably, inhibiting adolescent GABAergic neurons altered behavior in adolescents, but not adults (**Fig. 5e**). In adolescence, inhibiting GABAergic neurons initially reduced the acquisition of Hit and False Alarm trials, followed by an increase in the False Alarm acquisition rate on the second day of CNO administration. Adolescent lick rates to Hit, False Alarms and ITI showed a similar pattern (**Extended Data Fig. 7**), indicating that GABAergic inhibition could result in broad alterations in motivation. This may represent a scenario in which function is constricted as mice transition from adolescence into adulthood.

## Discussion

Our findings now provide various mechanisms that account for adolescent reward motivation (**Fig. 6**). At a behavioral level, we found that adolescent mice respond to more reward (CS+) and no-reward (CS-) cues than adults, indicating increased motivation in adolescence. However, increased lick rate to CS+ but not CS- cues in adolescence, suggests that adolescent mice have some understanding that the CS- is not rewarded. Thus, the increase in motivation is not generalized. When assessing neuronal activity, we found that the adolescent mPFC over-represents reward predictive cues (CS+), is hyperactive during rewarded trials, and under-represents non-reward predictive cues (CS-), when compared to adults. Furthermore, adolescent neurons generally decoded CS identity and rewarded trials at equal or greater levels than adults, demonstrating that the adolescent mPFC contains sufficient information to modulate reward behaviors. Demonstrating that the adolescent mPFC relies primarily on CS+ neurons to encode reward outcomes and CS identity. However, adolescents under-use CS- encoding neurons, which may result in an inability to encode differences between the non-rewarded trials (Miss, Correct Rejection, False Alarm). In general, our findings suggest that the adolescent mPFC focuses on representing rewarded trials and reward events, while the adult mPFC strives to represent all behavioral outcomes, not just rewarded ones. Thus, the observed age-related differences in reward behavior emerge as a result of differences in mPFC encoding properties and the mPFC microcircuitry.

**Figure 6:**
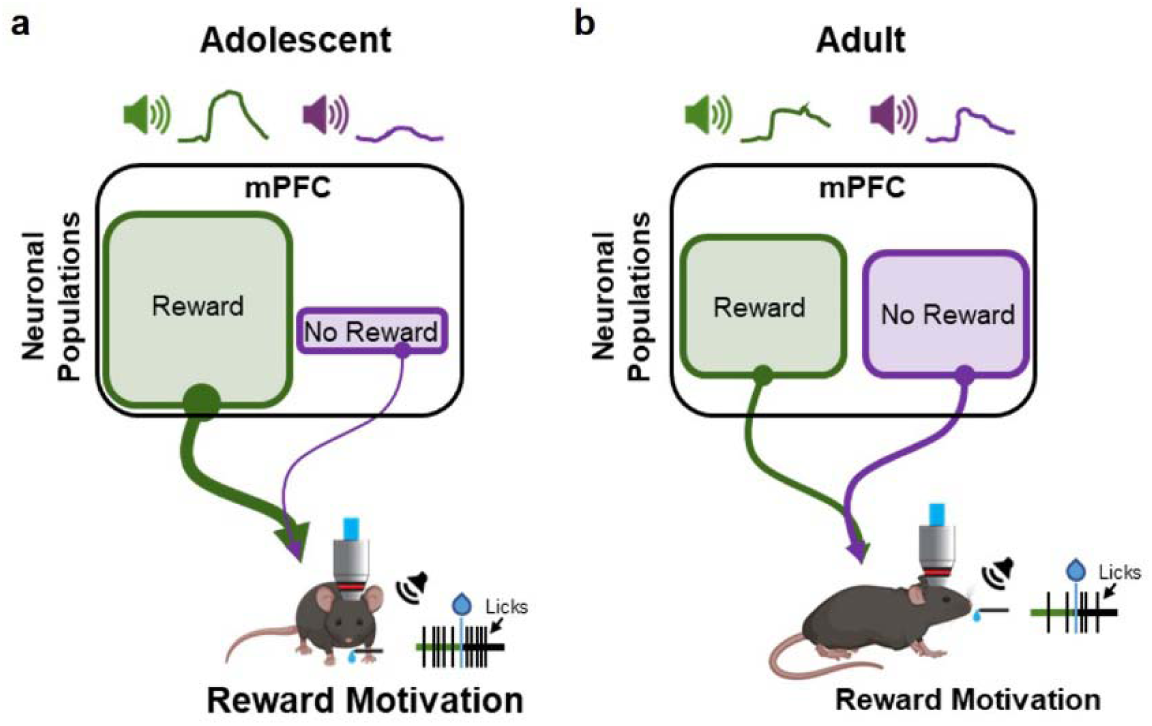
Proposed model for mPFC-driven increased adolescent reward motivation. a) Adolescent mice have an overrepresentation of CS+ encoding neurons, and an underrepresentation of CS- encoding neurons within the mPFC. This leads to increased reward-motivated behavior in adolescence, indexed by increased licking. b) In adults, the CS- encoding population is smaller than the CS+ encoding population, but much larger than the CS- encoding population in adolescents. This balance between CS+ and CS- encoding neurons allows the adult to better control when they will lick, decreasing overall lick rates and allowing them to withhold licks to non-rewarding cues.

Importantly, we demonstrated that causal manipulations of glutamatergic and GABAergic neurons in the mPFC have profound effects on reward behavior that are age-dependent. In adults, glutamatergic neuron inhibition decreased Hit and False Alarm acquisition rates, but this effect was not observed in adolescents. Inversely, GABAergic neuron inhibition had more profound effects in adolescents compared to adults. Specifically, adolescent GABAergic neuron inhibition impacted the number of Hit and False Alarm trials acquired. In the adult mPFC, GABAergic neurons exert their function on the microcircuit through inhibition and disinhibition. In adults, inhibition of parvalbumin interneurons disinhibits glutamatergic projection neurons to gate behavior ^41,42^. However, during adolescence, parvalbumin neurons are still maturing, increasing in both density and dendritic arborization, which possibly leads to differences in the disinhibition network ^24,43–46^. Additional work will be needed to fully understand how development impacts mPFC inhibition/disinhibition microcircuits. Furthermore, we found that excitation of glutamatergic neurons tended to increase Hit and False Alarm acquisition rates across days, regardless of age (**Extended Data Table 1**). The adolescent mPFC is more active during Hit trials compared to adults; it’s possible that chemogenetic inhibition in the adolescent glutamatergic neuronal signal is not sufficient to decrease activity to a threshold where behavioral effects are observable. Adolescent glutamatergic neurons may be more hyperactive but also harder to inhibit.

Crucially, findings show that the adolescent mPFC has different properties and ability to modulate behavior than the adult mPFC. We outline mechanisms that drive the increase in adolescent reward behavior. Future experiments must be targeted to understand how information flow to the mPFC contributes to the differences we observed. From previous work, we know that reward-related projections to the mPFC (e.g. from the ventral tegmental area) change from adolescence into adulthood ^7,28,29^. It is possible that as mPFC afferents mature, they relay different reward-related information, in both magnitude and temporal dimensions, to mPFC GABAergic and glutamatergic neurons. In combination with our current work, such studies would modify our working model of mPFC, suggesting that the adolescent mPFC receives more reward information, altering mPFC internal calculations to value the pursuit of rewards more highly than the adult mPFC.

Taken together, our study reinterprets mPFC function during development, demonstrating that the adolescent mPFC is not hypoactive; on the contrary, representations of reward-predictive cues and reward outcomes are amplified in the adolescent mPFC. Our findings refine our understanding of mPFC maturation, by confirming that top-down control is enhanced across development but that this change is probably not due to a lack of mPFC engagement in adolescence. Instead, over-representation and hyper-activity in response to reward cues in the mPFC may underlie deficits in inhibitory control and implicate the mPFC as a direct contributor to increased reward-seeking behavior in adolescence. This re-interpretation of mPFC development reinforces the view that functional development of brain regions must be evaluated with a wider lens. Development is not just turning up the “volume” of a brain region, gradually acquiring adult properties, but may be more akin to changing the radio station, exhibiting different properties. Furthermore, it pushes us to re-examine how drugs and treatments that affect behavior in adults, particularly those that affect mPFC, may be acting differently in adolescence.

## Figures

**Extended Data Figure 1:**
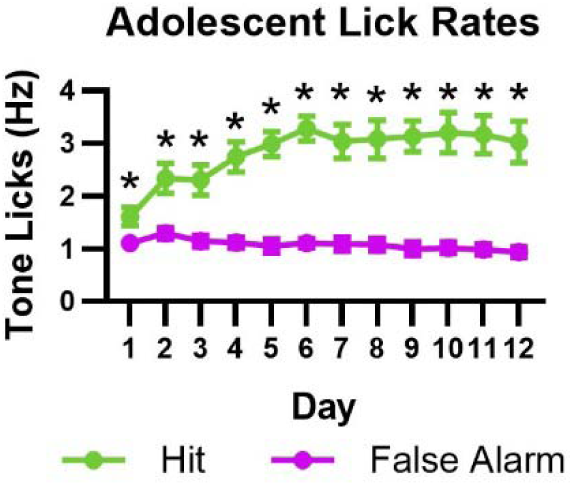
Adolescents have increased licking during Hit outcomes compared to False Alarm outcomes. Graph of lick rates to tones that resulted in a Hit or a False Alarm. A repeated measures two-way ANOVA revealed that adolescent mice had significant differences in the lick rate between trial outcomes (F_1,12_ = 106.4, p < 0.0001), and across days (F_3.27, 39.34_ = 2.90, p = 0.042), with a significant day by outcome type interaction (F_3.076, 36.91_ = 6.46, p = 0.0012). Šídák’s multiple comparison analysis comparing lick rates for each day found all days to be significantly different (p < 0.05). n=13 mice, the same mice are shown for Hit and False Alarm.

**Extended Data Figure 2:**
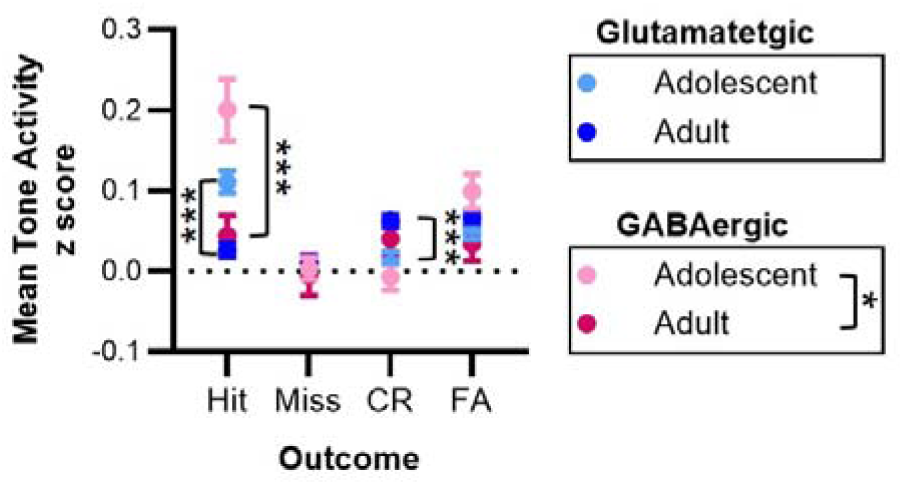
Adolescent neurons have increased activity during Hit outcomes. Analysis of the mean tone activity for adult and adolescent glutamatergic and GABAergic neurons. A three-way ANOVA for age, outcome, and cell type, revealed that age was a main source of variation (F_1,9792_ = 6.88, p = 0.0087). Significant main interaction of age by outcome (F _3, 9792_ = 16.14, p < 0.0001), age by cell type (F_1, 9792_ = 4.54, p = 0.033), and outcome by cell type (F_3, 9792_ = 3.63, p = 0.012) were also observed. A main effect of outcome was also observed (F_3, 9792_ = 16.62, p < 0.0001), indicating that neurons displayed differences in tone responses for the differing trial types. Follow-up two-way ANOVAs were conducted for each of the two neuronal subtypes (glutamatergic and GABAergic). In glutamatergic neurons, we found a main effect of outcome (F_3, 8563_ = 11.11, p < 0.0001) and an interaction between outcome and age (F_3, 8563_ = 21.48, p < 0.0001). Glutamatergic mean tone activity was significantly different between adolescent and adult mice during Hit (p < 0.0001) and correct rejection trials (p = 0.0009). For GABAergic neurons, a main effect of age (F 1, 1229 = 5.77; p = 0.016), outcome (F_3, 1229_ = 8.80; p < 0.0001), and interaction of age by outcome (F_3, 1229_ = 6.21; p = 0.0003) were detected. Mean tone activity during Hit trials were significantly different between adolescent and adult GABAergic neurons (p < 0.0001).

**Extended Data Figure 3:**
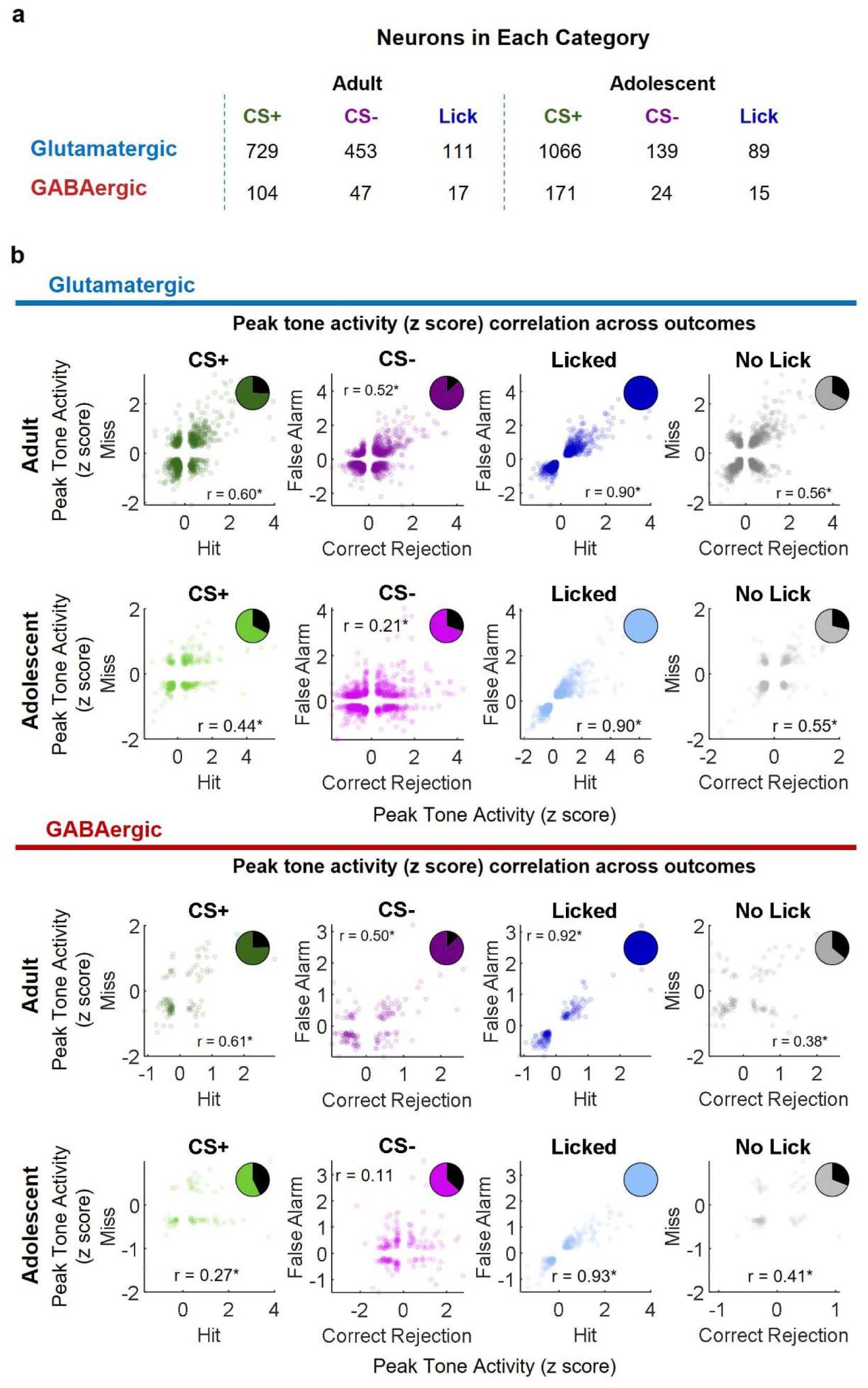
Stability of individual neurons across outcomes. a) Table showing the number of adult (left) and adolescent (right) neurons significant for each GLM predictor (CS+, CS-, Lick). Neurons significant for more than one predictor are represented more than once. b) Pairwise correlations of neuronal activity by trial outcome for each CS (CS+ and CS-), and behavior (Licked or No Lick). Each point represents the peak tone response magnitude (positive or negative) of a glutamatergic (top) or GABAergic (bottom) neuron in a given trial type. Only task-relevant neurons (GLM significant) are plotted. The linear coefficient of correlation (r) is presented on each graph, with * indicating a significant correlation (p < 0.05). Pie chart insets display the percent of neurons that showed a sign reversal in their peak response magnitude (black wedges), indicating a transition from inhibition (negative values) to excitation (positive values), or vice versa. The colored wedges represent the percent of neurons whose activity remained in the same direction (remained inhibited or remained excited) for both outcomes. Neurons exhibited sign reversal in their peak response magnitude (pie graph insets) in adult glutamatergic during CS+ (25.4%), CS- (13.1%), and No Lick (32.9%) trials and in adolescent glutamatergic neurons during CS+ (32.7%), CS- (30.3%) and No Lick (28.7%) trials. GABAergic neurons exhibited sign reversal in adults during CS+ (24.2%), CS- (12.9%), and No Lick (35.9%) trials and in adolescents during CS+ (42.8%), CS- (36.7%) and No Lick (30.9%) trials. No neuron exhibited sign reversal during Licked trials.

**Extended Data Figure 4:**
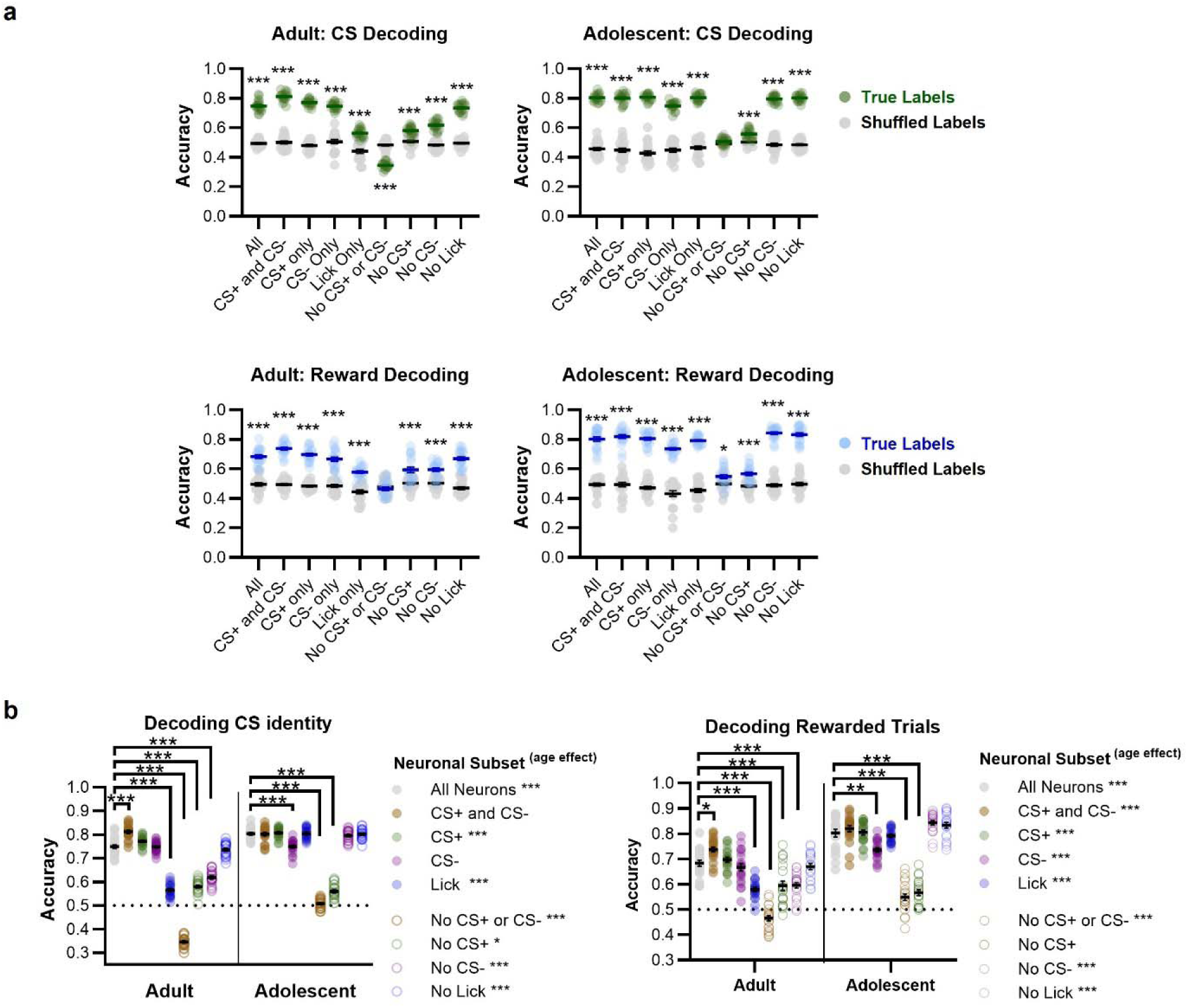
Neural decoding ability and within age comparisons. a) Graphs of the decoding accuracy for CS identity (top) and Rewarded trials (bottom), for adult (left) and adolescent (right) neuronal subsets compared to decoding accuracy of the same subsets trained on shuffled labels. Significance was tested using Šídák’s multiple comparison test. * p < 0.05, ** p < 0.01 *** p < 0.001. b) Decoding accuracy of CS identity (left) and Rewarded trials (Right) by distinct neuronal subpopulations. A Tukey multiple comparison analysis was used to determine which subsets performed significantly better or worse than the subset that contained all neurons (labeled All Neurons in the graph), and to determine significant differences in accuracy between adolescent and adult neuronal subsets (significance shown in figure legends). Individual values represent one decoding test. Individual runs and mean ± SEM for the decoder is shown. * p < 0.05, ** p < 0.01, *** p < 0.001

**Extended Data Figure 5:**
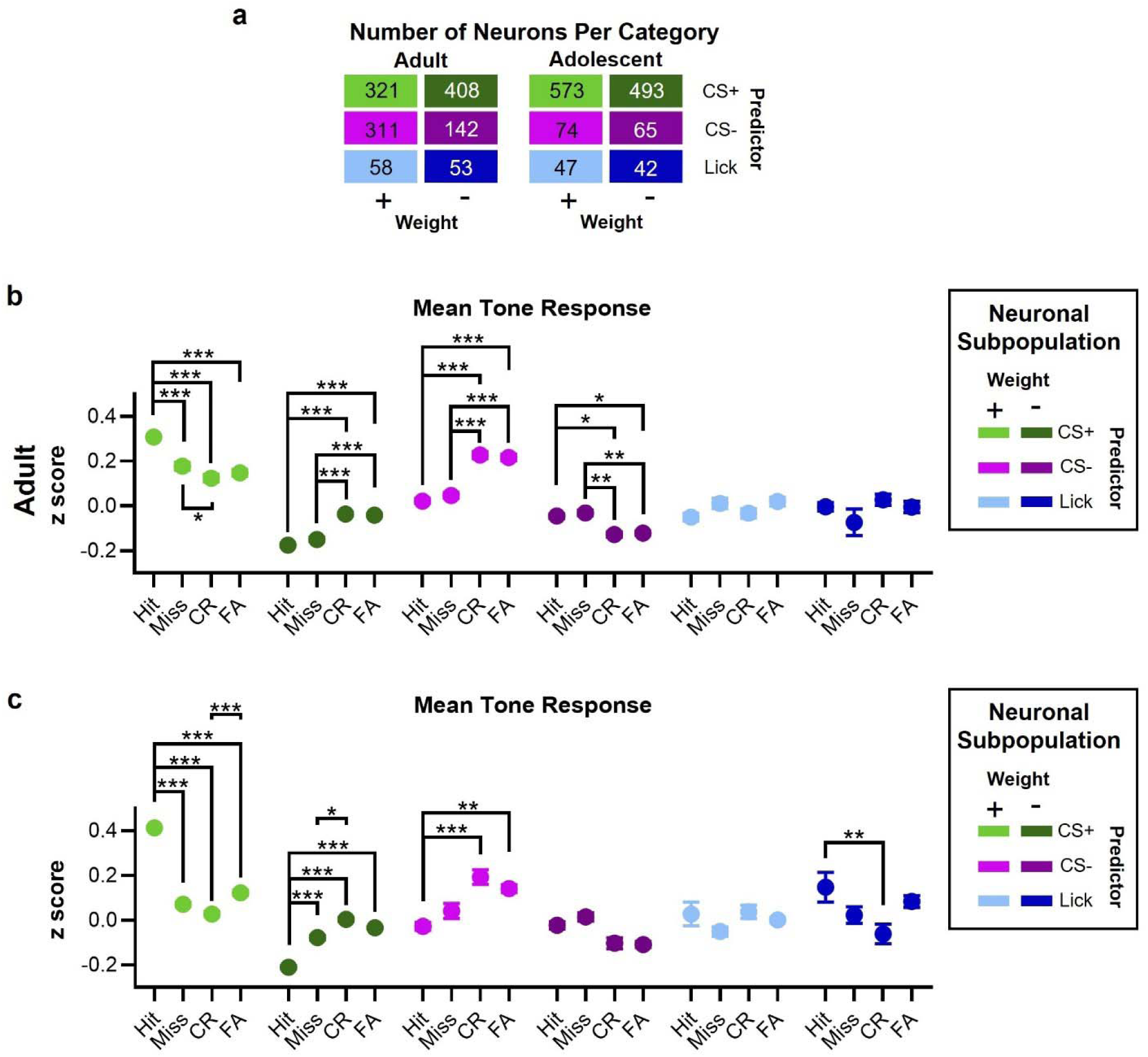
Adult CS- encoding neurons can distinguish between outcomes. a) Table containing the number of neurons in each category. b) Analysis of the mean tone activity for adult glutamatergic neuronal subpopulations across outcomes. A two-way ANOVA found a main effect of outcome (F_3, 5006_ = 4.51, p = 0.0036), subpopulation (F_5, 5006_ = 313.9, p < 0.0001) and an interaction between outcome and subpopulation (F_15, 5006_ = 29.37, p < 0.0001). A Tukey multiple comparison analysis was used to measure simple effects of mean tone response for each neuronal subpopulation across outcomes. For each comparison * p < 0.05, ** p < 0.01, *** p < 0.001. CR = Correct Rejection, FA = False Alarm. c) Analysis of the mean tone activity for adolescent glutamatergic neuronal subpopulations across outcomes. A two-way ANOVA found a main effect of subpopulation (F_5, 4247_ = 106.4, p < 0.0001) and an interaction between outcome and subpopulation (F_15, 4247_ = 51.33, p < 0.0001), but not outcome (F_3, 4247_ = 1.81, p = 0.14). A Tukey multiple comparison analysis was used to measure simple effects of mean tone response for each neuronal subpopulation across outcomes. For each comparison * p < 0.05, ** p < 0.01, *** p < 0.001. CR = Correct Rejection, FA = False Alarm.

**Extended Data Figure 6:**
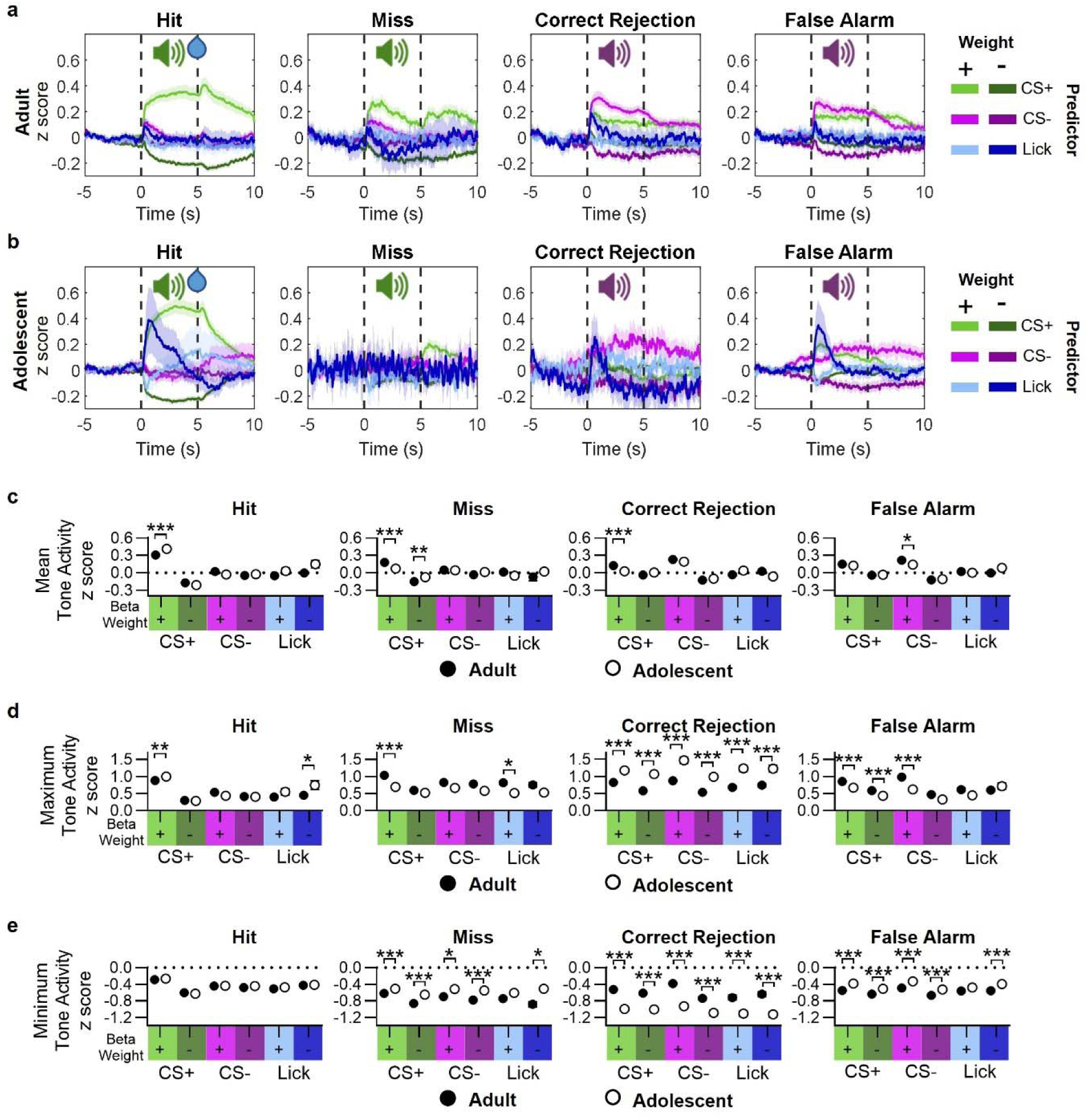
Neuronal subpopulations activity changes across outcomes. a) Plot of mean activity for adult neuronal subpopulations significant for each of the 3 predictors (CS+: green; CS-: magenta; and Lick: blue) during each outcome. Graphs show mean ± SEM (shaded area). b) Plot of mean activity for adolescent neuronal subpopulations significant for each of the 3 predictors (CS+: green; CS-: magenta; and Lick: blue) during each outcome. Graphs show mean ± SEM (shaded area). c) Comparison of mean tone activity of adult and adolescent neuronal subpopulations. Separate two-way ANOVA and Šídák test were run for each outcome. Two-way ANOVA was used to analyze each outcome for a main effect of age, neuronal subset, and interaction between age and neuronal subset. A main effect of age was only found for Hit (F_1, 2575_ = 5.694, p = 0.017). While a main effect of neuronal subset (Hit: F_5, 2575_ = 266.9, p < 0.0001; Miss: F_5, 1528_ = 57.54, p < 0.0001; Correct Rejection: F_5, 2575_ = 57.70, p < 0.0001; False Alarm: F_5, 2575_ = 102.1, p < 0.0001), and interaction between neuronal subset by age (Hit: F_5, 2575_ = 5.814, p < 0.0001; Miss: F_5, 1528_ = 8.791, p < 0.0001; Correct Rejection: F_5, 2575_ = 9.103, p < 0.0001; False Alarm: F_5, 2575_ = 2.893, p = 0.013) was found for all four outcomes. A Šídák’s multiple comparisons was used to determine simple effects of age, significance is shown on the graph. Mean ± SEM are shown. For each comparison * p < 0.05, ** p < 0.01, *** p < 0.001. d) Comparison of maximum tone activity of adult and adolescent neuronal subpopulations. Separate two-way ANOVA and Šídák test were run for each outcome. Two-way ANOVA was used to analyze each outcome for a main effect of age, neuronal subset, and interaction between age and neuronal subset. A significant main effect of age (Hit: F_1, 2575_ = 7.09, p = 0.0078; Miss: F_1, 1528_ = 28.06, p < 0.0001; Correct Rejection: F_1, 2575_ = 217.1, p < 0.0001; False Alarm: F _1, 2575_ = 37.40, p < 0.0001), neuronal subset (Hit: F_5, 2575_ = 172.2, p < 0.0001; Miss: F_5, 1528_ = 34.48, p < 0.0001; Correct Rejection: F_5, 2575_ = 22.41, p < 0.0001; False Alarm: F_5, 2575_ = 54.42, p < 0.0001), and interaction of neuronal subset by age (Hit: F_5, 2575_ = 4.46, p < 0.0005; Miss: F_5, 1528_ = 6.65, p < 0.0001; Correct Rejection: F_5, 2575_ = 2.39, p = 0.035; False Alarm: F_5, 2575_ = 5.15, p = 0.0001) was found in all four outcomes. A Šídák’s multiple comparisons was used to determine simple effects of age, significance is shown on the graph. Mean ± SEM are shown. For each simple comparison * p < 0.05, ** p < 0.01, *** p < 0.001. e) Comparison of maximum tone activity of adult and adolescent neuronal subpopulations. Separate two-way ANOVA and Šídák test were run for each outcome. Two-way ANOVA was used to analyze each outcome for a main effect of age, neuronal subset, and interaction between age and neuronal subset. A main effect of age was found for Miss (F_1, 1528_ = 49.97, p < 0.0001), Correct Rejection (F_1, 2575_ = 374.6, p < 0.0001), and False Alarm (F_1, 2575_ = 128.1, p < 0.0001) outcomes. A main effect of neuronal subset was found for each outcome (Hit: F_5, 2575_ = 295.7, p < 0.0001; Miss: F_5, 1528_ = 25.93, p < 0.0001; Correct Rejection: F_5, 2575_ = 15.18, p < 0.0001; False Alarm: F_5, 2575_ = 41.46, p < 0.0001). An interaction between neuronal subsets by age was observed for Miss (F_5, 1528_ = 2.44, p = 0.032) and Correct Rejection (F_5, 2575_ = 2.55, p = 0.025) outcomes. A Šídák’s multiple comparisons were used to determine simple effects of age, significance is shown on the graph. Mean ± SEM are shown. For each simple comparison * p < 0.05, ** p < 0.01, *** p < 0.001.

**Extended Data Figure 7:**
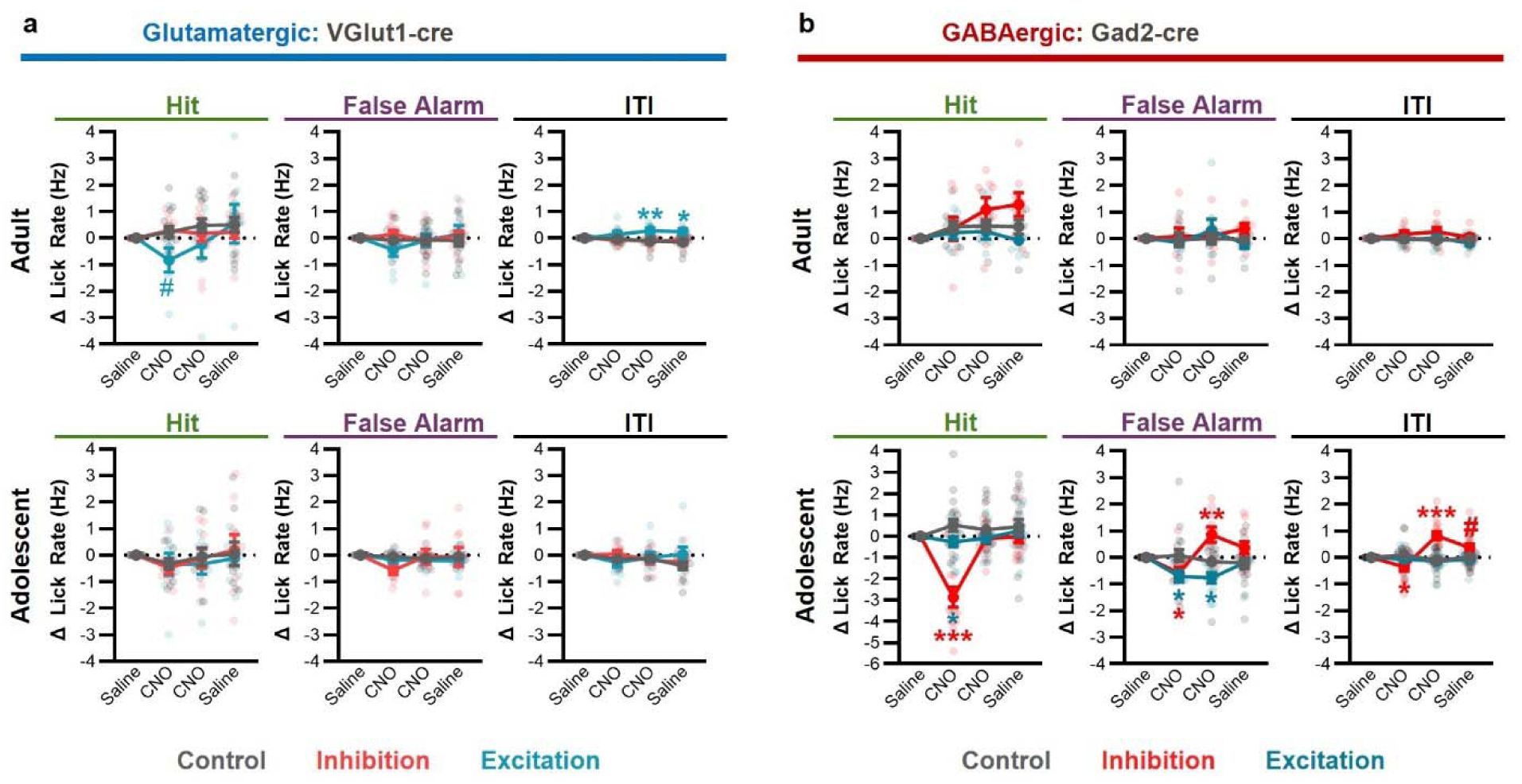
mPFC GABAergic neuron changes lick motivation in adolescents, but not adults. a) Inhibition (red) and excitation (blue) of glutamatergic mPFC neurons in adult (top) and adolescent mice. A Tukey’s multiple comparison analysis was run per graph. n = 8-14 mice per condition per age. # p <0.10, * p < 0.05, ** p < 0.01, *** p < 0.001 b) Inhibition (red) and excitation (blue) of GABAergic mPFC neurons in adult (top) and adolescent mice. A Tukey’s multiple comparison analysis was run per graph. n = 5-19 mice per condition per age. # p <0.10, * p < 0.05, ** p < 0.01, *** p < 0.001. For 3-way ANOVA analysis of age by treatment by day effects see **Extended Data Table. 1**

**Extended Data Table 1:**
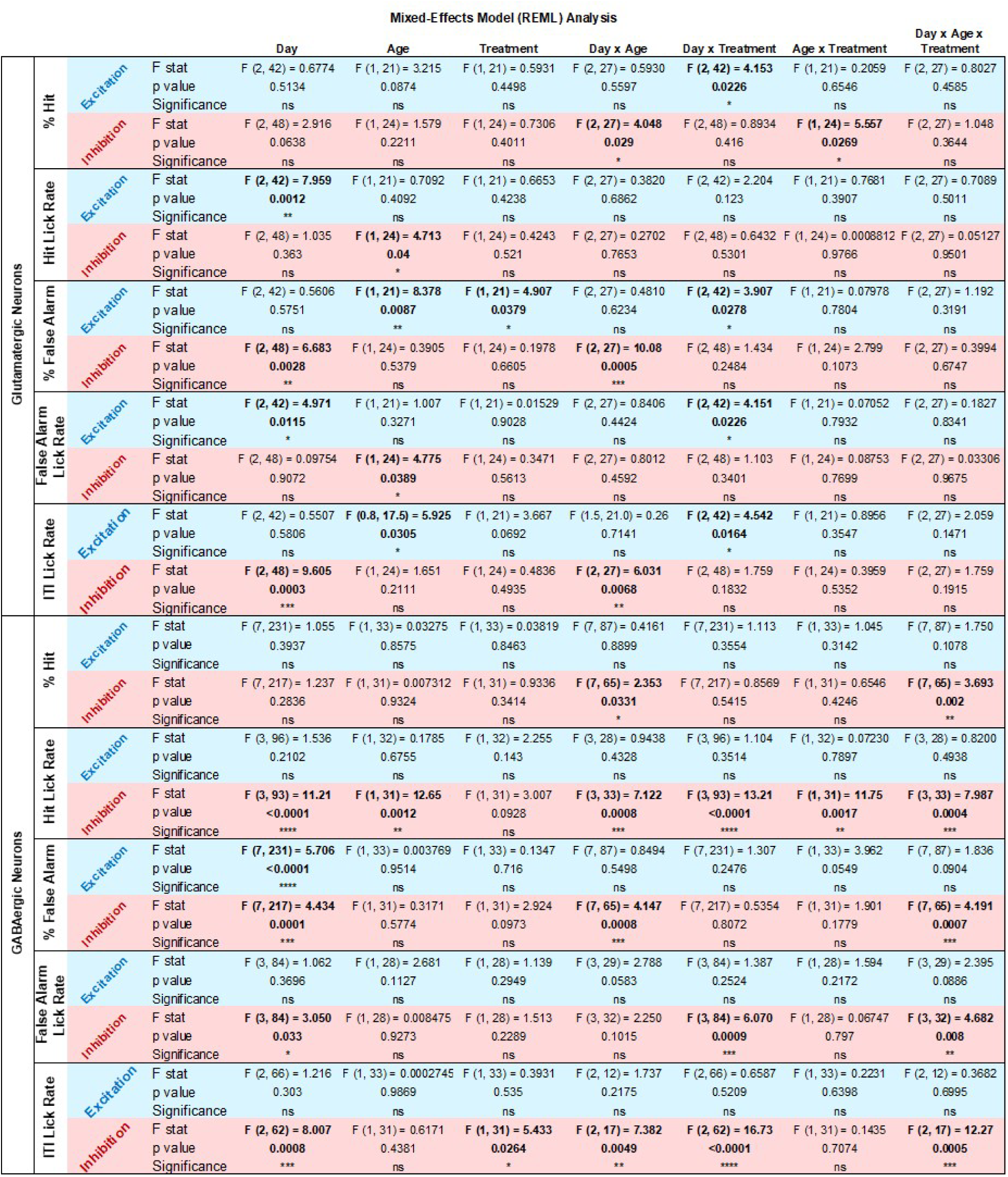
Statistical results for glutamatergic and GABAergic DREADD inhibition and excitation. Results of Mixed-Effects Model Analysis for age (adolescent versus adults), treatment (experimental versus control), and day (CNO day 1, CNO day 2, and Saline day 2).

## Materials and methods summary

### Subjects

Male and female C57BL/6 wildtype (Charles River Laboratories) were used for behavioral experiments. For 2-photon imaging Gad2-Cre (*Gad2tm2(cre)Zjh*/J, JAX#010802) ^47^ homozygous mice were crossed with Ai9 homozygous reporter mice (B6.Cg-*Gt(ROSA)26Sor^tm^*^9^*^(CAG-tdTomato)Hze^*/J, JAX#007909) ^48^ to produce heterozygous mice that expressed tdtomato in Gad2 (inhibitory) neurons. Heterozygous Gad2-Cre mice were used for chemogenetic inhibition of GABAergic neurons. Transgenic mouse lines were derived from a breeding stock acquired from Jackson Laboratories. All animals were housed according to NIH guidelines and maintained on a 12-hour light:dark cycle. Mice had free access to water and food except during experiments where mice underwent water restriction (see Active Reward Task section). Mice were weaned at P21, and group housed with littermates. Animal procedures are approved by Weill Cornell Medical College Institutional Animal Care and Use Committee, and consistent with the National Institutes of Health Guide for the Care and Use of Laboratory Animals.

### Behavioral Manipulations and Testing

#### General

For assessing developmental changes in neuronal activity during reward learning and expression we used a head-fixed modified pavlovian reward task (active reward task). Mice were trained through 3 phases: spout training (1 day, two 30 min. sessions), CS+ training (3 days, 1 session per day), and CS+/CS- training (7-12 days, 1 session per day). Training begins 7 days after head post-surgery, at P29 (± 1 day) for adolescence and at P75-P90 for adults. During spout training, mice are trained to lick sucrose (∼8uL of 10% sucrose) from a spout located directly in front of the mouse’s mouth. The reward spout is equipped with a touch sensor allowing for accurate detection and analysis of lick rates. As mice licked the spout during spout training day, the probability of acquiring a sucrose reward decrease. Spout training also served to habituate the mice to head fixation.

Having learned to lick the spout, mice then undergo CS+ reward conditioning. Mice receive 3 days of reward conditioning (CS+ only training session), consisting of 100 CS+ tone presentations (5 seconds duration, 70dB, 1000 or 9000 Hz), spaced out with a variable inter-trial-interval of 20-35 seconds. During these sessions mice had to lick the spout during the tone presentation to receive a sucrose droplet at tone offset. If mice did not lick the spout during the tone, they did not receive a reward at tone offset.

For CS+/CS- training, a CS- is introduced. The CS- is never rewarded. Depending on the experiment, mice are trained on CS+/CS- sessions for 7-12 days. CS+ / CS- conditioning consists of 50 random presentations of each of two distinguishable cues (5 seconds duration, 70dB, 1000 or 9000 hz), 100 tones total. A cue designates that the CS+ will predict sucrose reward, while the second cue, designated CS-, will predict no reward. When the CS+ is presented, mice must lick the spout to receive a reward at tone offset. There are four outcomes from this task: Hit (lick during CS+, rewarded), miss (no lick during CS+, not rewarded), False Alarm (lick during CS-, not rewarded), correct rejection (no lick during CS-, not rewarded).

#### Modifications of behavioral protocol for 2-photon imaging

2-photon imaging required modifications to the protocol to increase the number of trials acquired. Spout training was extended to 2 days with each day consisting of one session. This was done to allow mice to habituate to the noise generated by the 2-photon microscope. During CS+ only training sessions, 120 trials were presented, and the ITI was reduced to 10-15 seconds. During CS+/CS- training sessions, 120 trials were presented (60 CS+, 60 CS-), and the ITI was reduced to 10-15 seconds. The start of behavioral training was also delayed, starting at P30 for adolescents and P75 for adults, to allow additional time for prism clearing.

#### Modifications of chemogenetic inhibition and excitation

For chemogenetic inhibition. Mice were trained on four CS+/CS- sessions. On days 4 and 5 all mice were injected with saline (4ul saline / gram of mouse weight). On days 6 and 7 all mice were administered CNO (3 mg/kg) in saline (0.75 ug of CNO / ul of saline). On day 8 all mice were injected with saline (4ul saline / gram of mouse weight). Following behavioral experiments mice were perfused and brains were sliced for histological analysis to validate viral expression and injection site.

### Surgery

#### Head Fix Surgery

For head-fixed behavior, mice were implanted with a head-post. Adult (approx. P68) and adolescent (P22) mice were anesthetized with isoflurane gas anesthesia (2.0%–2.5% in 1 L/min oxygen), hair was shaved, and mice were placed in a stereotaxic apparatus. Buprenex (0.1 mg/kg, as an analgesic) was administered intraperitoneally. Tissue above the skull was cut away to reveal the skull, and lidocaine was applied. Skull was then cleaned with 0.3% hydrogen peroxide and left to dry (no more than 2 minutes). A custom-made head post was placed on top of the skull and secured with Metabond (Parkell Inc, Brentwood, NY). After the Metabond solidified, and implant was secure, isoflurane administration was stopped, and mice were allowed to wake. Mice are returned to their homecage and allowed at least 6 days to recover from surgery.

#### Prism Implant Surgery

For 2-photon imaging, surgeries were conducted in two stages to allow for sufficient viral expression. The first stage, viral injection, occurred between P1 and P3 for both adolescent and adult imaging groups. At P1-P3, neonates were placed in an isoflurane gas induction box (4% in 1 l/min oxygen), for at least 3 minutes. Anesthetized pups were then moved into a stereotaxic apparatus where isoflurane gas was administered at a lower concentration (2.0%–2.5% in 1 L/min oxygen). Skin was cleaned, and a small sliver (approximately. 1-2 mm) was cut above the mPFC. The central sinus was located, and an injector equipped with a 32-gauge beveled nanofil syringe (World Precision Instruments) preloaded with AAV1-syn-jGCaMP8f (Addgene: 162376-AAV1) was lowered to the skull level. The syringe was moved to the left of the midline and lowered 1.6 mm, where a 200 nL injection (75 nL/min.) was delivered. The injector was raised to 1.2 mm below the skull and a second 200 nL injection (75 nL/min.) was delivered. The syringe was then removed and a small amount of vetbond was applied to seal the cut on the pup’s skin. A second, more invasive surgery was conducted when mice reached P18 (for adolescent 2-photon imaging) or P63 (for adult 2-photon imaging). Mice were anesthetized in an isoflurane gas induction chamber (4% in 1 L/min oxygen), mouse hair was shaved, and mice were then moved to a stereotaxic apparatus where isoflurane gas was administered at a lower concentration (2.0%–2.5% in 1 L/min oxygen). Buprenex (0.1 mg/kg, analgesic) and Dexamethasone (2 mg/kg, anti-inflammatory) were co-administered intraperitoneally. Tissue above the skull was cut away to reveal the skull, and lidocaine was applied. Skull was then cleaned with 0.3% hydrogen peroxide. A ∼1.2 x 1.2 mm– diameter craniotomy was drilled rostral to the left mPFC. The dura was removed from above the surface of the brain to expose the tissue. A prism (1mm X 1mm X 2.5mm; M/L, A/P, D/V) was then inserted into the craniotomy to allow imaging of the ipsilateral mPFC. A small amount of Metabond (Parkell, Brentwood, NY) was used to secure the prism in place. A headpost was then placed around the prism and a second coat of Metabond was applied to secure the headplate and the prism implant. After the Metabond solidified, and implant was secure, isoflurane administration was stopped, and mice were allowed to wake. Dexamethasone was administered 24 hrs. and 48 hrs. post-surgery to decrease inflammation.

#### Chemogenetic Surgery

For fiber-photometry recordings of the mPFC, surgeries were conducted in two stages conducted at different ages. During the first surgery the virus was injected. During the second implants and headposts were secured. This allowed for sufficient viral expression (early surgery), and secure headposts (later surgery). The first surgery occurred at P16 (adolescent) and P59 - P70 (adults). Mice were anesthetized in an isoflurane gas induction chamber (4% in 1 l/min oxygen), mouse hair was shaved, and mice were then moved to a stereotaxic apparatus where isoflurane gas was administered at a lower concentration (2.0%–2.5% in 1 l/min oxygen).

Skin was cleaned, and the skin was cut along the midline. A borrow hole was drilled above mPFC (ML: ±0.32, AP: +1.5, DV: -2.1 for P16; ML: ±0.35, AP: +1.75, DV: -2.3 for adults). An injector equipped with a 32 gauge blunt nanofil syringe (preloaded with AAV5-hSyn-DIO-hM4D(Gi)-mCherry for neuronal inhibition, AAV5-hSyn-DIO-hM3D(Gq)-mCherry for neuronal excitation, or AAV-hSyn-DIO-mCherry for control; Addgene: 44362-AAV5, 44361-AAV5, 50459-AAV5) into mPFC and 200nL of virus was delivered. The syringe was then removed and vetbond was applied to seal the skin incision. For both age groups, a second surgery was conducted 6 days following the first surgery, to cement the headpost. Mice were anesthetized in an isoflurane gas induction chamber (4% in 1 L/min oxygen) and placed in a stereotaxic apparatus where isoflurane gas (2.0% - 2.5% in 1 L/min oxygen). Buprenex (0.1 mg/kg, analgesic) and Dexamethasone (2 mg/kg, anti-inflammatory) were co-administered intraperitoneally. Tissue above the skull was cut away to reveal the skull, and lidocaine was applied. Skull was then cleaned with 0.3% hydrogen peroxide. A headpost was then placed on the skull and Metabond was applied to cement the headpost. After the Metabond solidified, isoflurane administration was stopped, and mice were allowed to wake. Mice are returned to their homecage and allowed at least 6 days to recover from surgery.

### 2-photon imaging

2-photon calcium imaging was conducted using a commercial two-photon laser-scanning microscope (Olympus RS) equipped with a scanning galvanometer and a Spectra-Physics Mai Tai DeepSee laser tuned to 920nm, operating at 300-500mW. Single cells were recorded using a 10x 0.6NA water immersion objective, with 8mm working distance, over one plane. To minimize the impact of photobleaching, imaging was triggered by custom Matlab software, and acquired at 30 Hz starting 5 seconds before tone onset and ending 5 seconds after tone offset (15 seconds of continuous imaging), for all 120 trial presentations. Videos were analyzed using Suite2P analysis software ^49^. Cells were identified and visually inspected, and neuropil and calcium neuronal traces were exported. The neuropil signal was scaled and subtracted from the cell signal (FluorescenceCell - (.7 x FluorescenceNeuropil)). Each trial of each cell was normalized (z-scored) by the average activity of the same cell -5 to -2.5 seconds before tone onset. This allowed us to control for any bleaching that occurred during the session.

### General Linear Model

To identify task-relevant neurons, we used MATLAB 2023a’s *fitglm* to identify neurons that were significant for CS+, CS-, or lick variables. GLMs were modeled with normal distributions, as linear functions (no interaction effects) and with y-intercept terms. Each individual neuron was tested on a separate GLM. Each GLM was conducted on a neuronal activity trace that contained only pre-tone (5 sec) and tone (5 sec) activity acquired at 30hz. Given the goal of our study was to analyze mPFC’ s cue information, the reward period which produced activity in some cells was removed. If a predictor was determined to be significant for a given neuron, we obtained the F statistic (real F) comparing the fit of the full model to the fit of a reduced model (model without the significant predictor). To reduce the number of false positives, we then ran 500 iteration, where we circularly shifted (Matlab’s *circshift*) the neuronal traces and re-calculated the F statistic comparing the fit of the full model to the fit of a reduced model ^50^. Neurons with a significant p-value of the predictor (p<0.05) and whose real F statistic was greater than or equal to the 95th percentile of the shifted F statistic were considered to significantly encode the task feature. Any neuron that was significant for at least one of the 3 task-relevant predictors (CS+, CS-, or lick) was classified as a task-relevant neuron.

### Linear decoding with SVM classifiers

To test the ability of sub-populations to decode the identity of the CS or rewarded trial (**Fig. 3**), we used maximum-margin linear decoders (MATLAB *fitcsvm*). For each age, all neurons were pooled together into one matrix. Twenty iterations were run for each neuronal subpopulation. For CS decoding, at each iteration, the 120 CS trials were randomly separated into training (70%) and testing (30%) sets. For reward decoding, at each iteration, 25 rewarded and 25 unrewarded trials were randomly chosen from the 120 CS trials given, and separated into training (approx. 70%, rounded up) and testing (30%) sets. Only the 5 seconds of tone activity was used for decoding. The mean and SEM of the accuracy of the test set were plotted. To determine the random decoding accuracy of each subpopulation, we ran 20 iterations of training sets that had shuffled labels and tested the model on a test set with true values. Significance in decoding is determined by analyzing the significance of training with true labels compared to training with randomized labels.

### General Statistical Analysis

A two-tailed student’s t-test was used to compare between two groups. When more than two groups were assessed, the appropriate ANOVA was performed as stated in the figure legends. All ANOVA tests were followed by a multiple comparison test as detailed in the appropriate figure legend.

### Software

All ANOVA and multiple comparison tests were performed using Prism Graphpad statistical and graphing software. GLM, SVM, and correlation analysis were performed using Matlab (version 2023a), behavioral protocols and 2-P triggering were performed using Matlab (version 2022b). For all statistical analyses, statistical significance was defined as p<0.05. Data was graphed using Prism Graphpad and Matlab software. Schematics and illustrations were created using BioRender, Powerpoint, and Illustrator. ChatGPT was used for minimal editing.

### Data Availability

Upon publication, data analyzed for this work will be available at Weill Cornell’s Research Data Repository (RDR). Due to the large size, raw data will be available upon request.

### Code Availability

For this work, we used standard methods and functions readily available in Matlab (versions 2022b and 2023a). The behavioral task was coded within Matlab software and is available on GitHub https://github.com/GabaManzano/Reward-Tasks.

## Author Contributions

GMN developed experimental questions, designed and conducted experiments. SB, RR, and CL contributed to data interpretation and data visualization. RR conducted neuronal excitation and inhibition DREADD experiments with GMN. SB provided input and feedback on conceptual ideas, experimental design, and analysis pipelines. GMN wrote the initial manuscript, extensive edits on the initial manuscript were made by SB, RR, and CL.

## Acknowledgments

This work was supported by funding by the National Institutes of Health, from NIMH, and the Hope for Depression Research Foundation to CL, and from NIMH (MH124183; MH132082) to GMN. We would like to thank Dr. Hyeyoung Shin from Seoul National University for guidance on 2P video analysis, Dr. Arif Hamid from University of Minnesota for inspiring the design of our behavioral task, and Dr. Margaux Kenwood for feedback on decoder analysis.

## Notes

### Competing Interest Statement

The authors have declared no competing interest.

### Summary of Updates

Figures have been edited to facilitate direct comparisons between adolescent and adult findings. Causal experiments manipulating mPFC GABAergic and glutamatergic neurons have been added. Supplementary information is added.

